# Insect hormone PTTH regulates lifespan through temporal and spatial activation of NF-κB signaling during metamorphosis

**DOI:** 10.1101/2023.09.30.560323

**Authors:** Ping Kang, Peiduo Liu, Jinoh Kim, Ankur Kumar, Marie Bolton, Wren Murzyna, Zenessa J. Anderson, Lexi N. Frank, Nicholas Kavlock, Elizabeth Hoffman, Chad C. Martin, Marlene K. Dorneich-Hayes, Ting Miao, MaryJane Shimell, Weihang Chen, Yanhui Hu, Jo Anne Powell-Coffman, Michael B. O’Connor, Norbert Perrimon, Hua Bai

**Author notes:** **Corresponding Author:** Hua Bai, Ping Kang. **Author Contributions:** P.K. and H.B. designed research; P.K., P.L., J.K., A.K., M.B., W.M., Z.J.A., L.N.F., N.K., E.H., M.K.DH, T.M. performed research; M.S., M.B.O., W.C., Y.H. and N.P. contributed new reagents or analytic tools; P.K., P.L., C.C.M., W.C., Y.H. and H.B. analyzed data; P.K., J.A.P. and H.B. wrote the paper. **Competing Interest Statement:** The authors declare no competing interests.

## Abstract

The prothoracicotropic hormone (PTTH) is a well-known neuropeptide that regulates insect metamorphosis (the juvenile-to-adult transition) by inducing the biosynthesis of steroid hormones. However, the role of PTTH in adult physiology and longevity is largely unexplored. Here, we show that *Ptth* loss-of-function mutants are long-lived and exhibit increased resistance to oxidative stress in *Drosophila*. Intriguingly, we find that loss of *Ptth* blunt age-dependent upregulation of NF-κB signaling specifically in fly hepatocytes (oenocytes). We further show that oenocyte-specific overexpression of *Relish/NF-κB* blocks the lifespan extension of *Ptth* mutants, suggesting that PTTH regulates lifespan through oenocyte-specific NF-κB signaling. Surprisingly, adult-specific knockdown of *Ptth* did not prolong lifespan, indicating that PTTH controls longevity through developmental programs. Indeed, knockdown of PTTH receptor *Torso* in prothoracic gland (PG) during fly development prolongs lifespan. To uncover the developmental processes underlying PTTH-regulated lifespan, we perform a developmental transcriptomic analysis and identify an unexpected activation of NF-κB signaling in developing oenocytes during fly metamorphosis, which is blocked in *Ptth* mutants. Importantly, knockdown of *Relish/NF-κB* specifically in oenocytes during early pupal stages significantly prolongs the lifespan of adult flies. Thus, our findings uncover an unexpected role of PTTH in controlling adult lifespan through temporal and spatial activation of NF-κB signaling in developing hepatocytes and highlight the vital role of developmental NF-κB signaling in shaping adult physiology.

**Significance Statement:** Despite the strong link between animal development and adult lifespan, we know little about how developmental programs impact adult longevity, and when and where such programs are activated during development. Here, we demonstrate that loss of insect hormone PTTH prolongs lifespan and healthspan by repressing chronic inflammation in *Drosophila*. Intriguingly, we demonstrate that PTTH regulates adult lifespan through temporal and spatial activation of NF-κB signaling in developing hepatocytes during insect metamorphosis. These findings provide novel insights into the developmental programs that impact adult longevity.

## Introduction

Although aging is commonly viewed as a progressive deterioration of physiological function and accumulation of stochastic damage with age (1), there is evidence suggesting that the adult lifespan can be affected by changes occurring during development. Indeed, across animal species, both developmental duration and timing of sexual maturity positively correlate with adult lifespan (2, 3). In addition, mutations targeting either somatotropic axis in mammals (growth hormone (GH)/insulin-like growth factor (IGF) axis) or insulin/insulin-like growth factor signaling (IIS) in invertebrates often lead to retarded growth and prolonged adult lifespan (4–6). A number of previous lifespan studies in model organisms further support the existence of developmental programs as determinants of adult lifespan. For example, early-age treatment of GH reverses the lifespan extension of Ames dwarf mice (GH deficiency) (7). In addition, RNA interference-mediated knockdown of mitochondrial electron transport chain complex subunits throughout entire life, but not during adulthood, prolongs nematode lifespan (8). Similarly, the constitutive knockdown of complex I subunit NDUFS1/ND75 in muscles led to reduced systemic insulin signaling and extended lifespan in *Drosophila* (9). Further, transient exposure to low dosages of oxidants during larval development extends the adult lifespan of *Drosophila* (10). However, it remains unclear how exactly the developmental programs regulate adult longevity, and when and where such programs are activated during development.

In insects, particularly holometabolous insects, tissue growth and body size are achieved through multiple larval molts. At the end of larval development, animals undergo a unique process and transform into sexually mature adults during metamorphosis (11–13). Insect molting and metamorphosis are orchestrated by the steroid hormone ecdysone, which is synthesized and released from the prothoracic gland (PG) (14). Neuropeptide prothoracicotropic hormone (PTTH) is known as the major driver of ecdysone biosynthesis. PTTH is secreted by a few neuroendocrine cells in each brain hemisphere and signals through the receptor tyrosine kinase Torso to activate MAP kinase signaling within the PG (15, 16). PTTH belongs to the cystine knot family of growth factors (17). Although PTTH has no clear mammalian ortholog, it has been proposed that PTTH may play similar roles as mammalian gonadotropin-releasing hormone (GnRH) in controlling the timing of the juvenile-to-adult transition (18, 19). Loss of *Ptth* results in slower kinetics of ecdysone production, a delay in developmental timing, and slow imaginal disc growth (15, 20). However, the role of PTTH in insect metamorphosis (adult organ formation during the pupal stages) and adult physiology (e.g., lifespan) remains largely unexplored, despite the fact that *Ptth* transcript levels are much higher during pupal and adult stages than during larval stages (**Fig. 5C and S4**). Given the previously reported role of ecdysone signaling in longevity control in *Drosophila* (21, 22), it is reasonable to speculate that PTTH may also regulate adult lifespan beyond its developmental role.

Age-associated chronic inflammation, also known as inflammaging, is one of the major hallmarks of aging (1). Inflammatory cytokines, such as interleukin 6 (IL-6) and tumor necrosis factor α (TNF-α), are often induced during aging, and elevated IL-6 in the circulation is a powerful indicator of all-cause mortality in aging human populations (23, 24). Chronic inflammation is not only a biomarker of aging, but also drives aging and age-related pathologies. Anti-inflammatory interventions often preserve tissue function and slow aging processes. For example, inhibition of TNF-α signaling rescues premature aging phenotypes in mice with Tfam-deficient T cells (25). Further, brain-specific knockout of *IKKβ* in mice (26). As in the mammalian system, insect innate immunity, in particular the immune deficiency pathway (Imd), is the first line of defense against bacteria, fungi, and other pathogens (27). Glial-specific knockdown of *Relish/NF-κB* in *Drosophila* prolongs lifespan (28). Upon infection, the transcription factor Relish/NF-κB is activated through a conserved signal transduction cascade involved in peptidoglycan recognition proteins (PGRPs), Imd/RIP1 kinase, caspase Dredd, TGF-β activated kinase 1 (TAK1), and IκB kinase complex (IKK) (29). Relish positively regulates the expression of antimicrobial peptide genes (AMPs), such as Diptericin DptA, a group of small peptides with unique inhibitory effects against pathogens.

In *Drosophila*, ecdysone signaling primes innate immunity and Relish/NF-κB signaling through the transcriptional control of peptidoglycan recognition protein LC (PGRP-LC) in *Drosophila* (30, 31), possibly linking developmental hormonal signaling and innate immunity. During the larva-to-pupa transition in holometabolous insects, there is a large pulse of ecdysone that initiates the prepupal stage and the shutdown of larval wandering behavior (32, 33), suggesting that the prepupal pulse of ecdysone primes the Relish/NF-κB pathway to facilitate the immune defense at the immobile pupal stage. However, the role of innate immunity during insect metamorphosis is largely unexplored. Recently, Relish/NF-κB has been found to be expressed in the hematopoietic niche during larval development in *Drosophila*, and to play a vital role in maintaining the blood progenitors in developing lymph glands (34). Intriguingly, silencing Relish specifically at the *Drosophila* pupal stage enhances the susceptibility of adult flies to viral infection, indicating that Relish/NF-κB signaling during metamorphosis is essential in conditioning adult antiviral responses (35). These studies suggest that NF-κB signaling could be a novel developmental program that controls adult longevity through remodeling insect metamorphosis.

In this study, we examined the role of PTTH in the regulation of longevity and Relish/NF-κB signaling. We show that loss of *Drosophila Ptth* extends lifespan and enhances resistance to oxidative stress in both males and females. The lifespan extension of *Ptth* loss-of-function (LOF) mutants is dependent on age-dependent activation of Relish/NF-κB, especially in the fly hepatocytes (oenocytes). Intriguingly, we found that Relish/NF-κB signaling is activated in oenocytes during *Drosophila* pupal development, and this temporal and spatial activation of NF-κB signaling is blocked by the loss of *Ptth*. Intriguingly, oenocyte- and pupal-specific knockdown of *Relish/NF-κB* significantly extends lifespan. Taken together, our findings uncover an unexpected role of PTTH in controlling adult lifespan through temporal and spatial activation of NF-κB and provide novel insights into the developmental programs that impact adult longevity.

## Results

### Loss of *Ptth* prolongs lifespan and healthspan in *Drosophila*

To determine the role of PTTH in longevity regulation, we utilized three previously generated loss-of-function alleles of *Ptth* (20, 36). Two of them, *Ptth^8K1J^* (6 bp deletion in the final exon) and *Ptth^120F2A^* (7 bp deletion in the final exon), were previously generated through TALEN-directed mutagenesis (20) (**Fig. S1A**). Before setting up the lifespan analysis, we backcrossed these two *Ptth* alleles to wild-type background (*w^1118^*) for 5 generations to eliminate the confounding effects of genetic background. In addition, we backcrossed the third allele, a knockout mutation of *Ptth* (36), which we named *Ptth^TI^*, to wild-type background (y*w^R^*) for five generations. *Ptth^TI^* mutants are likely a null allele, as all the exons were deleted and replaced by a 3P3-RFP cassette through CRISPR-Cas9 and homologous recombination-mediated gene targeting (36) (**Fig. S1A**).

As in previous studies (20), all backcrossed *Ptth* mutants showed delayed pupariation (**Fig. S1B and S1C**). Interestingly, these slow-growing *Ptth* mutants were all long-lived compared to their match controls (both females and males, **Fig. 1A-1D**), and showed reduced age-specific mortality (**Fig. S1D**). The lifespan extension of *Ptth^8K1J^* females was relatively smaller (**Fig. 1A**), suggesting that *Ptth^8K1J^* is a weak allele, likely due to small amino acid changes. Consistent with the extended lifespan, *Ptth* mutants exhibited increased resistance to paraquat-induced oxidative stress (both females and males, **Fig. 1E and 1F**), and preserved climbing ability during aging (both females and males, **Fig. 1G and 1H**). Importantly, *Ptth* mutants prolong their lifespan without any reproductive cost. In fact, the female fecundity of *Ptth* mutants was higher than that of wild-type flies (**Fig. S2A**). Altogether, our data demonstrate that PTTH regulates lifespan and healthspan in *Drosophila*, beyond its developmental role.

**Fig 1.**
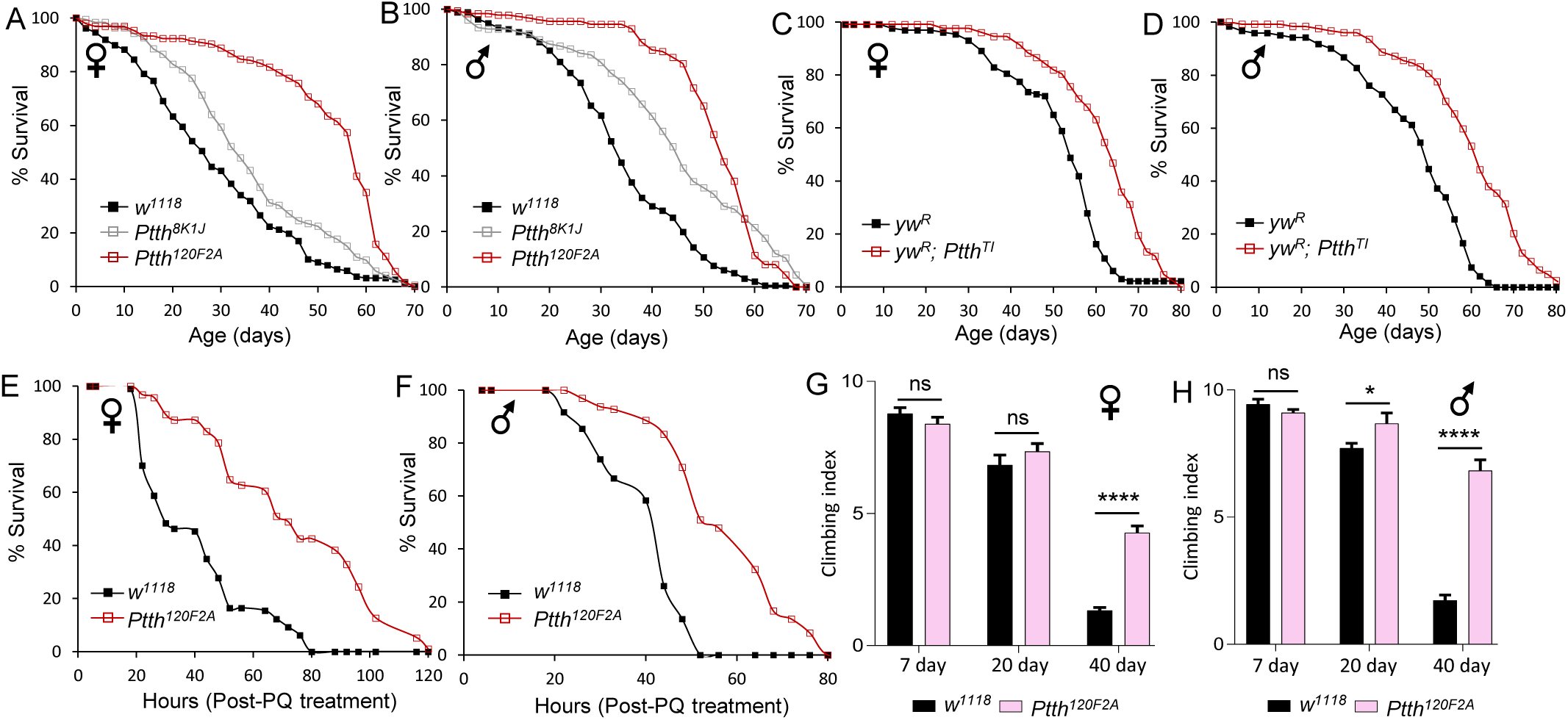
Loss of *Ptth* prolongs lifespan and healthspan in *Drosophila*. (*A* and *B*) Lifespan analysis of two loss-of-function alleles of *Ptth*. Log-rank test (vs. *w^1118^*). *Ptth^8K1J^* (Female: *p <* 0.05, n = 192. Male: *p* < 0.001, n = 182); *Ptth^120F2A^* (Female: *p* < 0.001, n = 197. Male: *p* < 0.001, n = 184). *Ptth^8K1J^* is a weak allele, likely due to small amino acid changes. (*C* and *D*) Lifespan analysis of loss-of-function allele of *Ptth^TI^*. Log-rank test (vs. *yw^R^*). Female: *p <* 0.05, n = 128. Male: *p* < 0.001, n = 123. (*E* and *F*) Survival analysis of *Ptth* mutants under 20 mM paraquat (PQ) treatment in both males and females. (log-rank test (vs. *w^1118^*), *p* < 0.001, n = 100). (*G* and *H*) Climbing activity of female and male *Ptth* mutants during aging. Two-way ANOVA followed by Bonferroni’s multiple comparison test. ns, not significant; * *p* < 0.05; **** *p* < 0.0001. n = 30.

### *Ptth* mutants repress age-dependent upregulation of innate immunity signaling

To understand the molecular mechanisms underlying PTTH-regulated longevity, we performed a bulk RNA-Seq analysis to characterize the transcriptomic changes in young (5-day-old) and aged (38-day-old) female wild-type (*w^1118^*) and *Ptth* mutants (*Ptth^120F2A^*, an allele with strong lifespan extension). There were 731 differentially expressed genes (DEGs) between *Ptth* mutants and wild-type at young age (fold change > 1.5, FDR < 0.05) (**Fig. 2A**). Among them, 129 were significantly upregulated, while 602 were significantly downregulated. Gene ontology (GO) analysis showed that these genes are enriched in biological processes, such as digestive system development, mesoderm development, epithelial tube morphogenesis, and tissue morphogenesis and development (**Fig. 2B**). On the other hand, there were 610 DEGs between *Ptth* mutants and wild-type at old age (fold change > 1.5, FDR < 0.05) (**Fig. 2C**). Surprisingly, almost all the enriched pathways identified through GO analysis are related to innate immunity (**Fig. 2D**).

**Fig 2.**
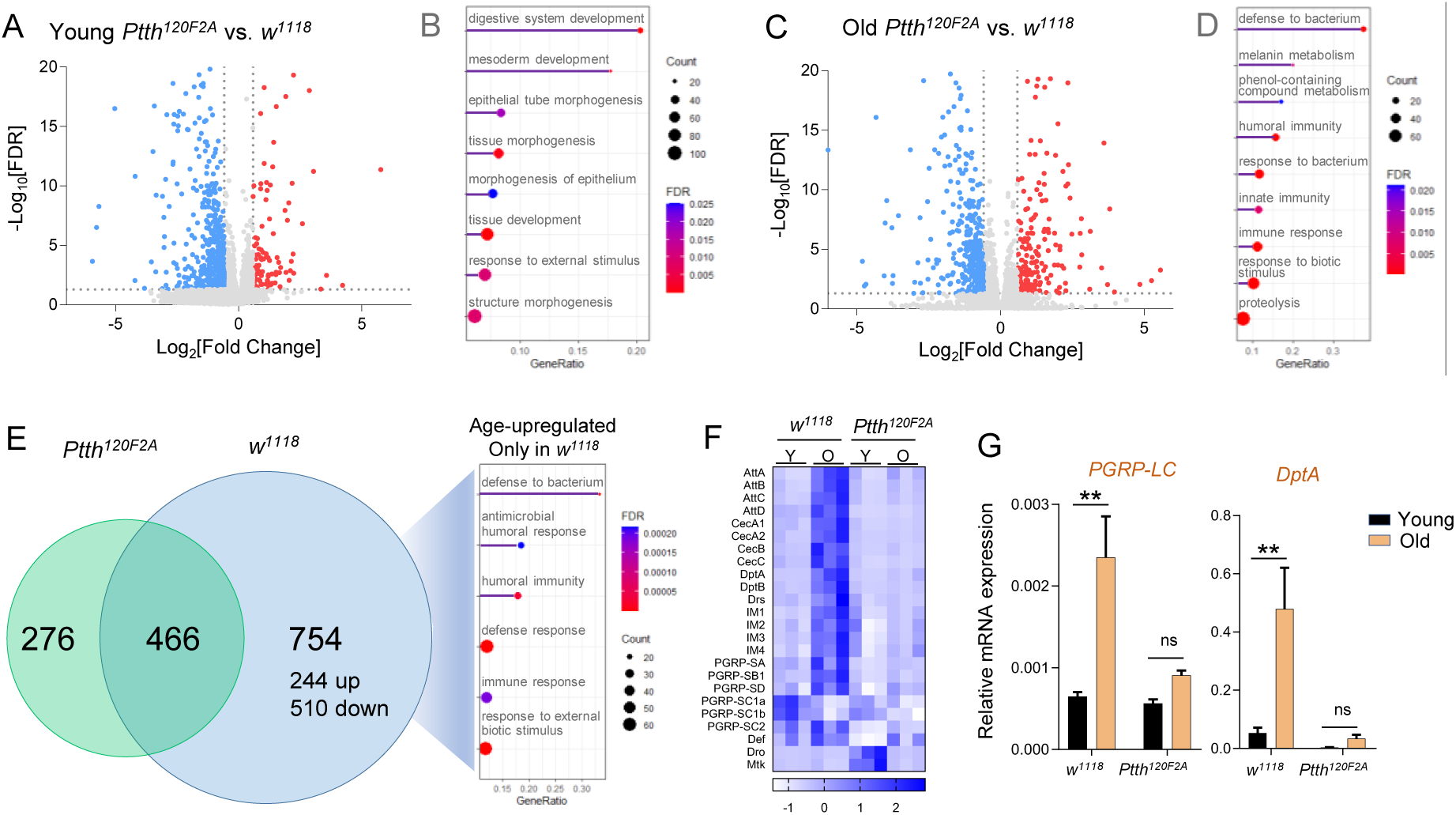
*Ptth* mutants repress age-dependent upregulation of innate immunity signaling. (*A* and *C*) Volcano plot showing genes significantly upregulated and downregulated by *Ptth* mutants at young (5-day-old) and old age (38-day-old). Fold change > 1.5, FDR < 0.05. (*B* and *D*) Dot plot analysis showing differentially regulated biological processes between *Ptth* mutants and wild-type (*w^1118^*) at young and old ages. (*E*) Venn diagram and dot plot showing the number of age-upregulated genes in *Ptth* mutants and wild-type (*w^1118^*), and the upregulated biological processes only found in wild-type (*w^1118^*). (*F*) Heat map showing genes in innate immunity pathway are differentially regulated by *Ptth* mutants with age. Y: young age; O: old age. (*G*) qPCR analysis of the expression of peptidoglycan recognition protein LC (*PGRP-LC*) and antimicrobial peptide Diptericin A (*DptA*) in young and old *Ptth* mutants and wild-type (*w^1118^*). Two-way ANOVA followed by Bonferroni’s multiple comparison test. ns, not significant; ** *p* < 0.01. n = 3.

To gain insights into the biological processes induced in aged wild-type flies but not in the *Ptth* mutants, we analyzed the age-associated DEGs in both wild-type and *Ptth* mutants, respectively. Among the 1220 age-associated DEGs found in wild-type flies (fold change > 2, FDR < 0.05), 754 (244 upregulated and 510 downregulated) were differentially expressed only in aged wild-type flies, but not in aged *Ptth* mutants (**Fig. 2E**). GO analysis revealed that the 244 age-associated DEGs found in wild-type were again enriched for immune response and defense to bacterium (**Fig. 2E**). As shown in the heatmap (**Fig. 2F**), most of the AMP genes (e.g., *AttA, AttB, AttC, CecA1, CecA2, CecB, CecC, DptA, DptB, Drs*) and Bomanin genes (e.g., *IM1/BomS1, IM2/BomS2, IM3/BomS3, IM4/Dso1*) were significantly induced in aged wild-type flies, but not in *Ptth* mutants. Thus, loss of *Ptth* blocks the age-dependent induction of both Imd and Toll innate immunity pathways. Further, we verified these findings using qRT-PCR. The expression of both *PGRP-LC* and *DptA*, two major players of the Imd pathway, was significantly upregulated upon normal aging in wild-type flies, while loss of *Ptth* alleviated these age-related inductions (**Fig. 2G**). As the hyperactivation of innate immune pathways is a hallmark of chronic inflammation (inflammaging), it suggests that PTTH regulates lifespan through innate immunity, and that reduced PTTH signaling suppresses inflammaging.

To test whether innate immunity in *Ptth* mutants could result in any cost of animal fitness, in particular in fighting bacterial infections, we challenged young wild-type and *Ptth* mutant females with the Gram-negative pathogenic bacterium *Erwinia carotovora carotovora 15* (*Ecc15*). Surprisingly, *Ptth* mutants were more tolerant to *Ecc15* infection than wild-type flies (**Fig. S2B**). *Ecc15* challenge upregulated innate immunity (elevated transcription of *PGRP-LC* and *DptA*) in both wild-type and *Ptth* mutants, even though the degree of induction was much lower in *Ptth* mutants (**Fig. S2C**). In addition, we also examined the inflammatory signaling, such as JAK-STAT signaling, in *Ptth* mutants upon *Ecc15* challenge. Interestingly, *Ptth* mutants significantly attenuated *Ecc15*-induced JAK-STAT signaling, indicated by the expression of *upd3* (homolog of mammalian IL-6) and *Socs36E* (**Fig. S2D**). It is known that JAK-STAT signaling is activated in response to gut epithelial cell damage during bacterial infection and plays an essential role in intestinal repair (37). The low levels of JAK-STAT activation upon *Ecc15* challenge indicates that loss of *Ptth* protects gut epithelial cells from damage through optimal levels of immune response and reduced chronic inflammation during bacterial infection. The reduced chronic inflammation, indicated by the expression of *upd3/IL-6* and *Socs36E*, was also found in aged *Ptth* mutants when compared to wild-type flies (**Fig. S2E**). Together, these results demonstrate that *Ptth* mutants exhibit robust immune defense capacity with well-balanced innate immunity activation and reduced chronic inflammation during bacterial infection and aging.

### PTTH regulates Relish/NF-κB signaling specifically in fly hepatocytes

PTTH is a neuropeptide hormone secreted by two bilateral small populations of neuroendocrine cells (PG neurons) in the larval brain. It can act as a neurotransmitter that is transported from PG neurons to PG to promote ecdysone biosynthesis (15). On the other hand, it behaves as a systemic hormone that travels through the circulation to target distal tissues (e.g., light-sensing organs) to modulate larval light avoidance behavior (38). However, how PTTH regulates adult physiology is largely unknown. To identify the target adult tissues through which PTTH regulates longevity and innate immunity (in particular the Imd pathway), we monitored the age-dependent induction of *DptA* in fly tissues. Strikingly, we noticed that among dissected heads, thorax and abdomen from female adult flies, loss of *Ptth* blocked age-dependent induction of *DptA* only in fly abdomen (data not shown).

Next, we examined three major abdominal tissues of female adults (fat body, oenocytes, and gut) for Relish immunostaining and *DptA* mRNA expression to monitor age-related changes in innate immunity signaling in wild-type and *Ptth* mutants. Nuclear translocation of Relish, the key transcription factor of the Imd pathway, is known as the hallmark of innate immunity activation (27, 39) that regulates the expression of AMP genes (e.g., *PGRP-LC* and *DptA*). Upon activation of innate immune response (such as aging and bacterial infection), Relish is cleaved by rapid proteolytic cleavage, resulting in a 68 kDa N-terminal fragment (Rel68) and a 49 kDa C-terminal fragment (Rel49). Rel49 is degraded in the cytoplasm while Rel68 translocates to the nucleus to activate the transcription of AMP genes (e.g., *DptA*) (40). Using an antibody specifically recognizing Rel68, we found that nuclear localized Relish was detected in the fat body, but not oenocytes and midgut, in young wild-type and *Ptth* mutants (**Fig. 3A**). During aging, nuclear translocation of Relish was enhanced in the fat body and midgut (seen in non-enterocytes) in both wild-type and *Ptth* mutants (**Fig. 3A**). Interestingly, the age-dependent induction of nuclear translocation of Relish was only observed in wild-type oenocytes, but not in *Ptth* mutant oenocytes (**Fig. 3A**). Consistently, we found that loss of *Ptth* blocked the age-related induction of *DptA* expression only in oenocytes, but not in the fat body and gut (**Fig. 3B**). Further, age-dependent activation of JAK-STAT signaling, as indicated by the expression of *upd3/IL-6* and *Socs36E*, was also blocked by *Ptth* mutants specifically in oenocytes (**Fig. S3A and S3B**). Taken together, these findings suggest that PTTH regulates age-dependent activation of innate immunity and inflammation specifically in fly oenocytes, the homolog of mammalian hepatocytes.

**Fig 3.**
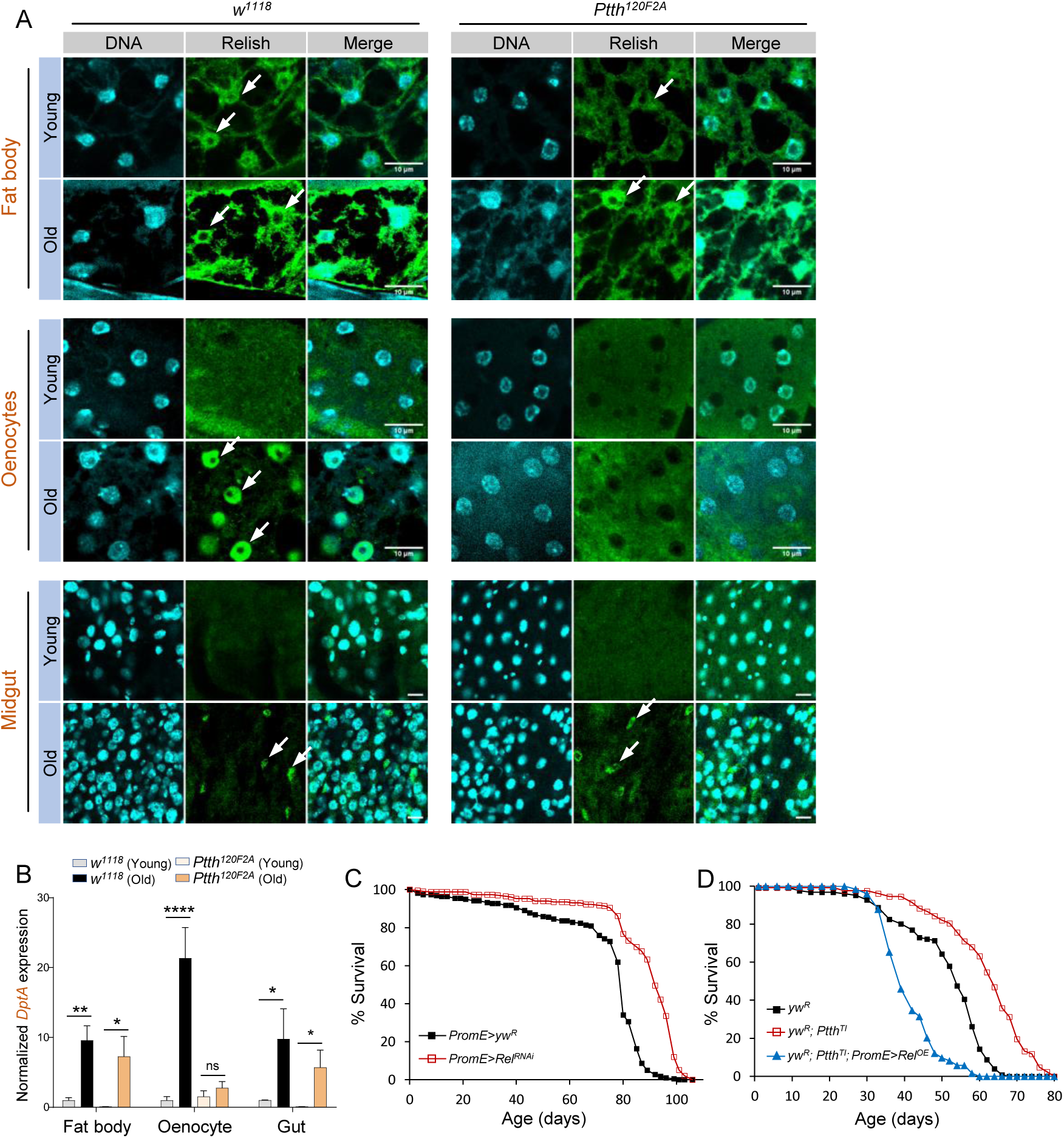
Loss of *Ptth* blocks age-dependent activation of NF-κB signaling specifically in fly hepatocytes (oenocytes). (*A*) Immunostaining analysis of nuclear translocation of Relish/NF-κB in young and old *Ptth* mutants and wild-type (*w^1118^*). Three tissues were analyzed, oenocytes, midgut, fat body. Scale bar: 10 μm. White arrow: nuclear localized Relish. (*B*) qPCR analysis of the expression of Diptericin A (*DptA*) in three different fly tissues dissected from young and old *Ptth* mutants and wild-type (*w^1118^*). Two-way ANOVA followed by Bonferroni’s multiple comparison test. ns, not significant; * *p* < 0.05; ** *p* < 0.01; **** *p* < 0.0001. n = 3∼6. (*C*) Oenocyte-specific knockdown of *Relish/NF-κB* extends lifespan (female). Log-rank test, *p* < 0.001, total n = 452. (*D*) Oenocyte-specific overexpression of *Relish/NF-κB* blocked the lifespan extension of *Ptth* mutants (female). Log-Rank test (*Ptth^TI^* vs. *Ptth^TI^ ; PromE>Rel^OE^*), *p* < 0.001, total n = 378.

### Hepatic Relish/NF-κB is required for the lifespan extension of *Ptth* mutants

NF-κB signaling has been shown to regulate longevity through the central nervous system. Brain-specific knockout of *IKKβ* in mice (26) or glial-specific knockdown of *Relish/NF-κB* in flies (28) prolongs lifespan. However, it remains to be determined whether hepatic NF-κB signaling also contributes to longevity. Thus, we knocked down *Relish* specifically in oenocytes using the oenocyte-specific GAL4 driver (*PromE-GAL4*). Strikingly, oenocyte-specific knockdown of *Relish* extended the lifespan of female flies (**Fig. 3C**), and reduced age-specific mortality (**Fig. S3C**).

Given that PTTH regulates age-dependent nuclear translocation of Relish in adult oenocytes, we tested whether the lifespan extension of *Ptth* mutants is dependent on oenocyte-specific Relish/NF-κB signaling. We combined *Ptth* mutants with the oenocyte-specific GAL4 driver (*PromE-GAL4*), then crossed it with a *UAS-FLAG-Rel.68* line to overexpress a constitutively active form of *Relish* in oenocytes in the *Ptth* mutant background. Interestingly, oenocyte-specific expression of Rel68 blocked the lifespan extension effects of *Ptth* mutants (**Fig. 3D and S3D**). Together, our data suggests that PTTH regulates lifespan through oenocyte-specific Relish/NF-κB signaling.

### PTTH regulates lifespan throughout the development

PTTH is secreted from PG neurons during larval development which degenerate during the pupal stage. These PTTH-positive neurons undergo developmental pruning and rewiring to form adult PTTH neurons (15). Interestingly, *Ptth* is expressed highly in prepupal and pupal stages, and relatively low expression of *Ptth* is detected in adult females (**Fig. S4**). To determine whether PTTH regulates lifespan in adults by targeting adult tissues (like oenocytes), we performed lifespan analysis of flies with adult-onset global knockdown of *Ptth* using a ubiquitous GeneSwitch driver (*Da-GS-GAL4*). Unexpectedly, adult-onset knockdown of *Ptth* shortened the lifespan of female flies (**Fig. 4A**), suggesting that PTTH might regulate lifespan during development, rather than during the adult stage.

**Fig 4.**
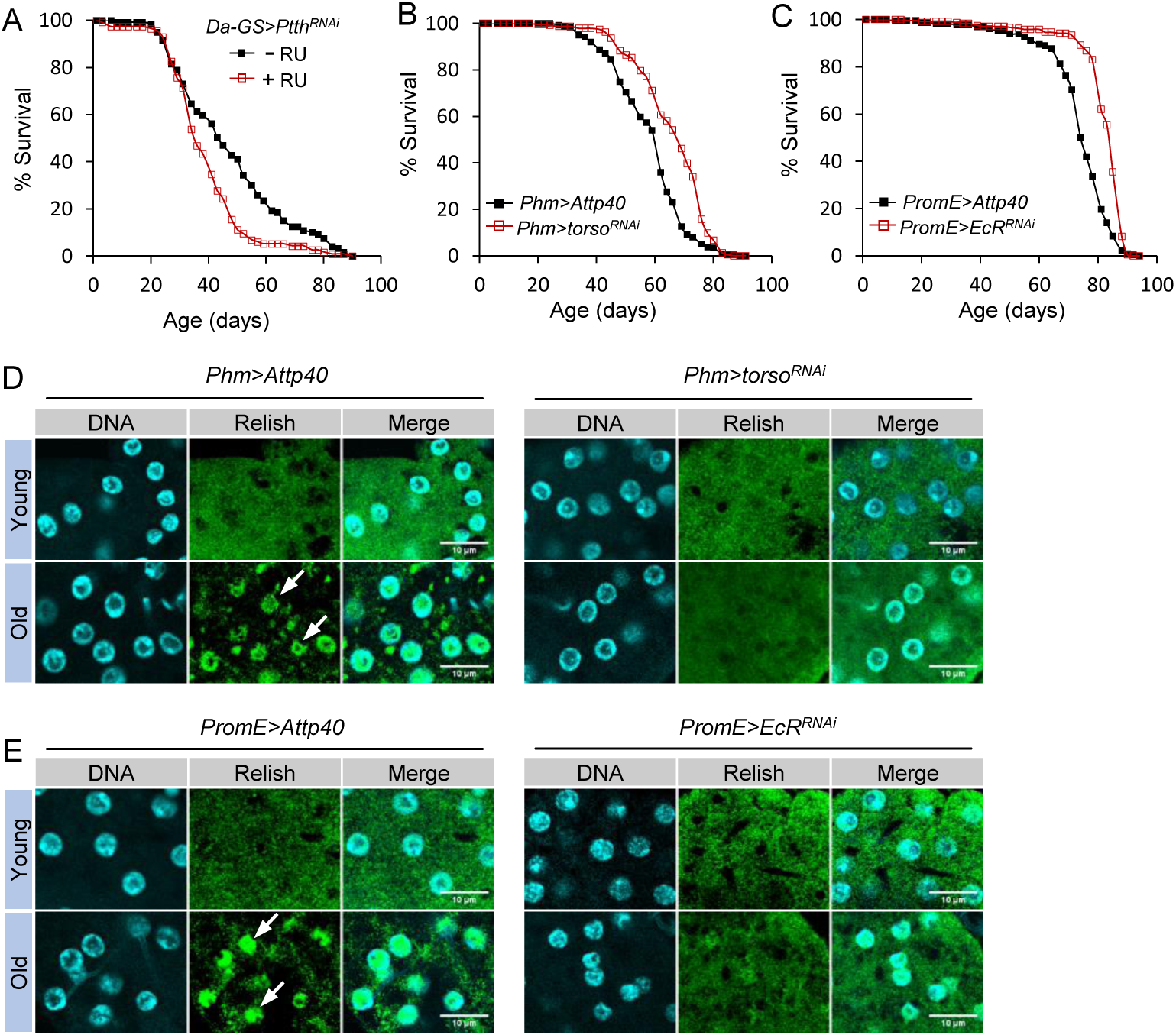
PTTH regulates lifespan through development. (*A*)) Lifespan analysis of adult-onset knockdown of *Ptth* (female). Log-rank test, *p* < 0.001, n = 234. RU486 (mifepristone, or RU) was used to activate Da-GS-GAL4 GeneSwitch driver. (*B*) Prothoracic gland (PG)-specific knockdown of PTTH receptor *Torso* extends lifespan (female). Log-rank test, *p* < 0.001, n = 475. (*C*) Oenocyte-specific knockdown of ecdysone receptor (*EcR*) extends lifespan (female). Log-rank test, *p* < 0.001, n = 471. (*D*) Immunostaining analysis of nuclear translocation of Relish/NF-κB in oenocytes of young and old control (*Phm>Attp40*) and PG-specific *Torso* knockdown flies (*Phm>torso^RNAi^*). Scale bar: 10 μm. White arrow: nuclear localized Relish. (*E*) Immunostaining analysis of nuclear translocation of Relish/NF-κB in oenocytes of young and old control (*PromE>Attp40*) and PG-specific *Torso* knockdown flies (*PromE>EcR^RNAi^*). Scale bar: 10 μm. White arrow: nuclear localized Relish.

If PTTH regulates lifespan through a developmental program, we wondered whether this is mediated through the PTTH receptor Torso in the PG during metamorphosis. The receptor tyrosine kinase Torso mediates PTTH signaling by activating a MAP kinase cascade within the PG to initiate ecdysone biosynthesis during *Drosophila* metamorphosis (16, 20). Using a PG-specific GAL4 driver (*Phm-GAL4*), we constitutively knocked down *Torso* in the PG and found that PG-specific knockdown of *Torso* prolonged the lifespan of female flies (**Fig. 4B**). Interestingly, PG-specific knockdown of *Torso* blocked age-dependent increases in oenocyte-specific nuclear translocation of Relish (**Fig. 4D**). Altogether, these data suggest that PTTH regulates lifespan and oenocyte-specific Relish/NF-κB signaling by targeting its receptor Torso in the PG during development and metamorphosis.

In the PG, Torso controls the production of ecdysone, which in turn promotes larval and pupal molts, the development of adult structures (16, 32), and innate immunity (31). To test whether ecdysteroid hormones activate ecdysone receptor (EcR) in oenocytes to modulate oenocyte-specific Relish/NF-κB signaling, and eventually lifespan, we performed a lifespan analysis of flies with oenocyte-specific knockdown of *EcR*. As expected, oenocyte-specific knockdown of *EcR* prolonged the lifespan of female flies (**Fig. 4C**). Consistently, oenocyte-specific knockdown of *EcR* also blunted age-dependent increases in nuclear translocation of Relish in oenocytes (**Fig. 4E**).

Taken together, our data support the model that PTTH binds to its receptor Torso in the PG to control the production and release of ecdysone during metamorphosis. Ecdysone then travels to oenocytes and activates EcR signaling to modulate adult oenocyte function, which in turn results in extended lifespan and protects oenocytes by lowering NF-κB activation and chronic inflammation during aging.

### Relish/NF-κB signaling is activated in developing oenocytes during *Drosophila* metamorphosis, which is blocked by *Ptth* mutants

To uncover the developmental processes through which PTTH signaling regulates lifespan and NF-κB signaling, we performed bulk RNA-seq to profile the transcriptomic changes throughout larva-to-adult development in both wild-type (*w^1118^*) and *Ptth* mutants (*Ptth^120F2A^*). Eight developmental stages were used: L3E (3^rd^ instar larvae, 48 hr prior to pupariation), L3L (3^rd^ instar larvae, 24 hr prior to pupariation), WP (white prepupa), P1 (one day post pupariation), P2 (two days post pupariation), P3 (three days post pupariation), P4 (four days post pupariation), A0 (1∼3 hr after adult eclosion). Principal component analysis (PCA) revealed that the biological replicates for each developmental stage grouped together, while the samples between groups were dispersed (**Fig. 5A**). The eight developmental groups, regardless of wild-type or *Ptth* mutants, were arranged perfectly following the developmental trajectory from L3E to A0 (**Fig. 5A**). In addition, there was a clear separation between wild-type and *Ptth* mutants in the two 3^rd^ instar larval stages (L3E and L3L), as well as the three pupal stages (P1, P2, P3) (**Fig. 5A**).

**Fig 5.**
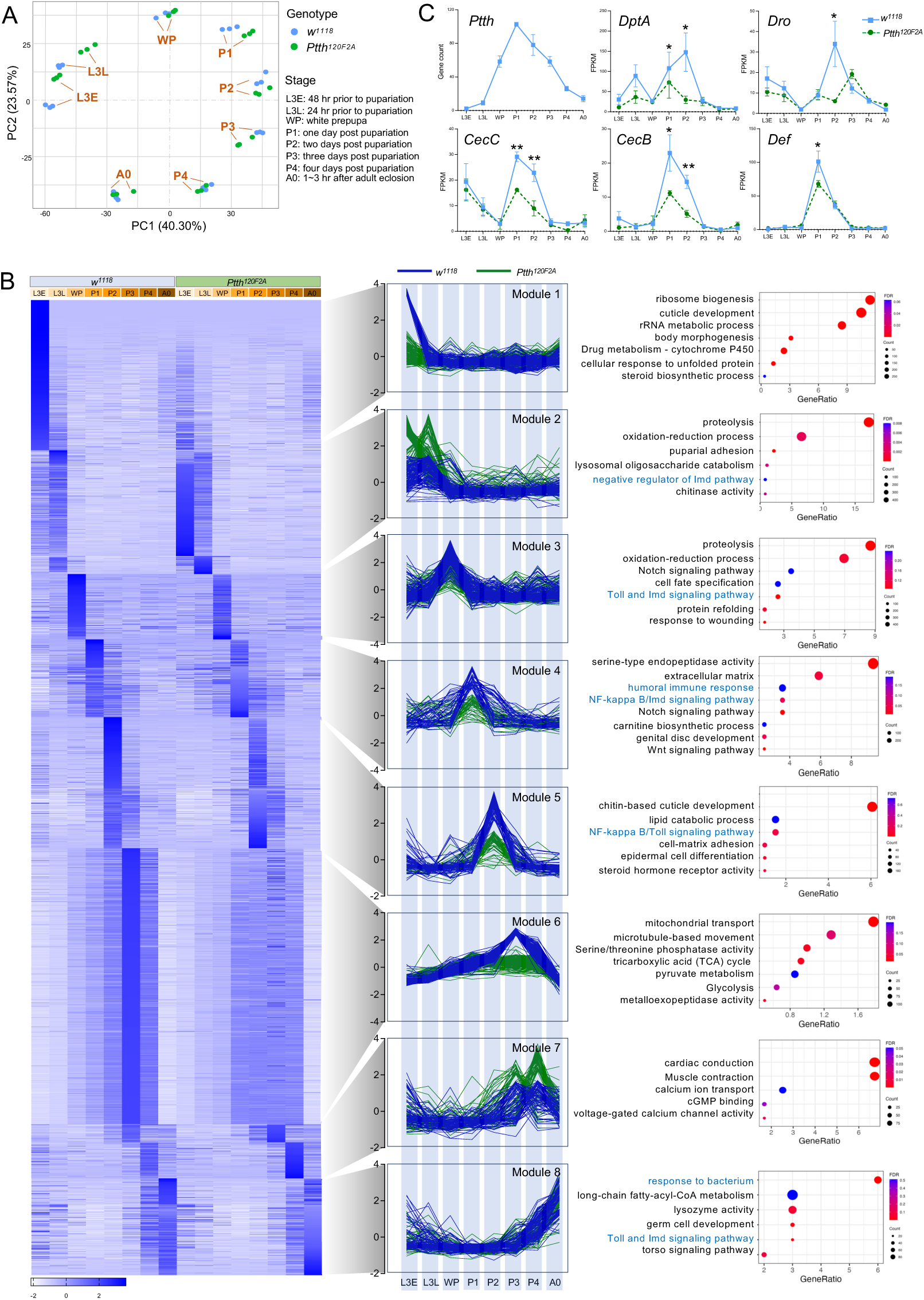
NF-κB signaling is activated during *Drosophila* metamorphosis, which is blocked by *Ptth* mutants. (*A*) PCA plot showing stage-specific transcriptomic profiling of wild-type (*w^1118^*) and *Ptth* mutants (*Ptth^120F2A^*). (*B*) Heat map, line plots, and pathways analysis for 8 distinct clusters identified from 4000 DEGs between wild-type (*w^1118^*) and Ptth mutants (*Ptth^120F2A^*) at different developmental stages. (*C*) Expression of *Ptth* gene and antimicrobial peptide genes in wild-type (*w^1118^*) and *Ptth* mutants (*Ptth^120F2A^*) at different developmental stages. * *p* < 0.05; ** *p* < 0.01. n = 3.

Although *Ptth* transcript levels peaked during the pupal stages (P1-P2) (**Fig. 5C**), more genes were differentially regulated by *Ptth* mutants at the two larval stages (**Fig. S5A**). As expected, the differentially regulated biological processes enriched for each developmental stage were also distinct. During larval development (L3E and L3L), cytoplasmic translation and ribosome biogenesis were differentially regulated by *Ptth* mutants. During pupal development, various metabolic and developmental processes were differentially regulated by *Ptth* mutants, such as cell differentiation, cell morphogenesis, epithelium development, and nervous system development (**Fig. S5C**).

To further characterize the biological processes that were enriched for specific developmental stages and specific genotypes, we performed DESeq2 differential expression analyses (41) to identify stage-specific genes for each genotype respectively (fold change > 2, FDR < 0.05), followed by DEG identification by comparing wild-type samples and *Ptth* mutant samples at each developmental stage (see method section). We identified a total of 4313 DEGs that were differentially expressed between wild-type and *Ptth* mutants across eight different developmental stages (**Fig. S5A**). Our analysis also revealed eight distinct co-expression gene modules (**Fig. 5B**). Each module represented a cluster of genes highly co-expressed at one specific developmental stage in wild-type samples. For example, Module 1 includes genes that were specifically induced at L3E stage in wild-type samples, whereas Module 3 includes genes that were specifically induced at WP stage in wild-type samples (**Fig. 5B**). Among the eight gene modules, loss of *Ptth* resulted in a decreased gene expression in most of the modules, except for Module 2 (L3L) and Module 7 (P3-P4) (**Fig. 5B**). Strong reduction of gene expression by *Ptth* mutants was observed in Module 1 (L3E) and Module 6 (P3). Module 1 includes genes involved in ribosome biogenesis, cuticle development, rRNA metabolic process, and body morphogenesis, whereas Module 6 includes genes in mitochondrial transport, microtubule-based movement, serine/threonine phosphatase activity, tricarboxylic acid (TCA) cycle (**Fig. 5B**). In contrast, loss of *Ptth* resulted in an increased gene expression in Module 2 (L3L) and Module 7 (P3-P4). Similar results were observed using multiWGCNA gene co-expression analysis (**Fig. S5B**).

Strikingly, both the innate immunity pathway and NF-κB signaling were found differentially regulated by *Ptth* mutants in five out of eight modules (Module 2, 3, 4, 5, 8) (**Fig. 5B**), suggesting a strong link between PTTH and NF-κB signaling during metamorphosis. Interestingly, most AMP genes (e.g., *DptA, CecB, CecC*) showed peak expression during early pupal stages (P1-P2), which corresponds to the peak of *Ptth* expression during metamorphosis (**Fig. 5C**), which is consistent with the significant reduction of AMP gene expression during early pupal stages associated with loss of *Ptth* (**Fig. 5C**).

Next, we monitored the nuclear translocation of Relish/NF-κB in oenocytes dissected from one-day-old pupae. Pupal tissues were co-stained with streptavidin to locate oenocytes (42). Consistent with our RNA-seq data, a strong nuclear translocation of Relish/NF-κB was observed in oenocytes dissected from one-day-old pupae, where Relish nuclear translocation was blocked by *Ptth* mutants (**Fig. 6A**). Altogether, we demonstrate that PTTH signaling is required for the activation of NF-κB signaling in developing oenocytes during metamorphosis.

**Fig 6.**
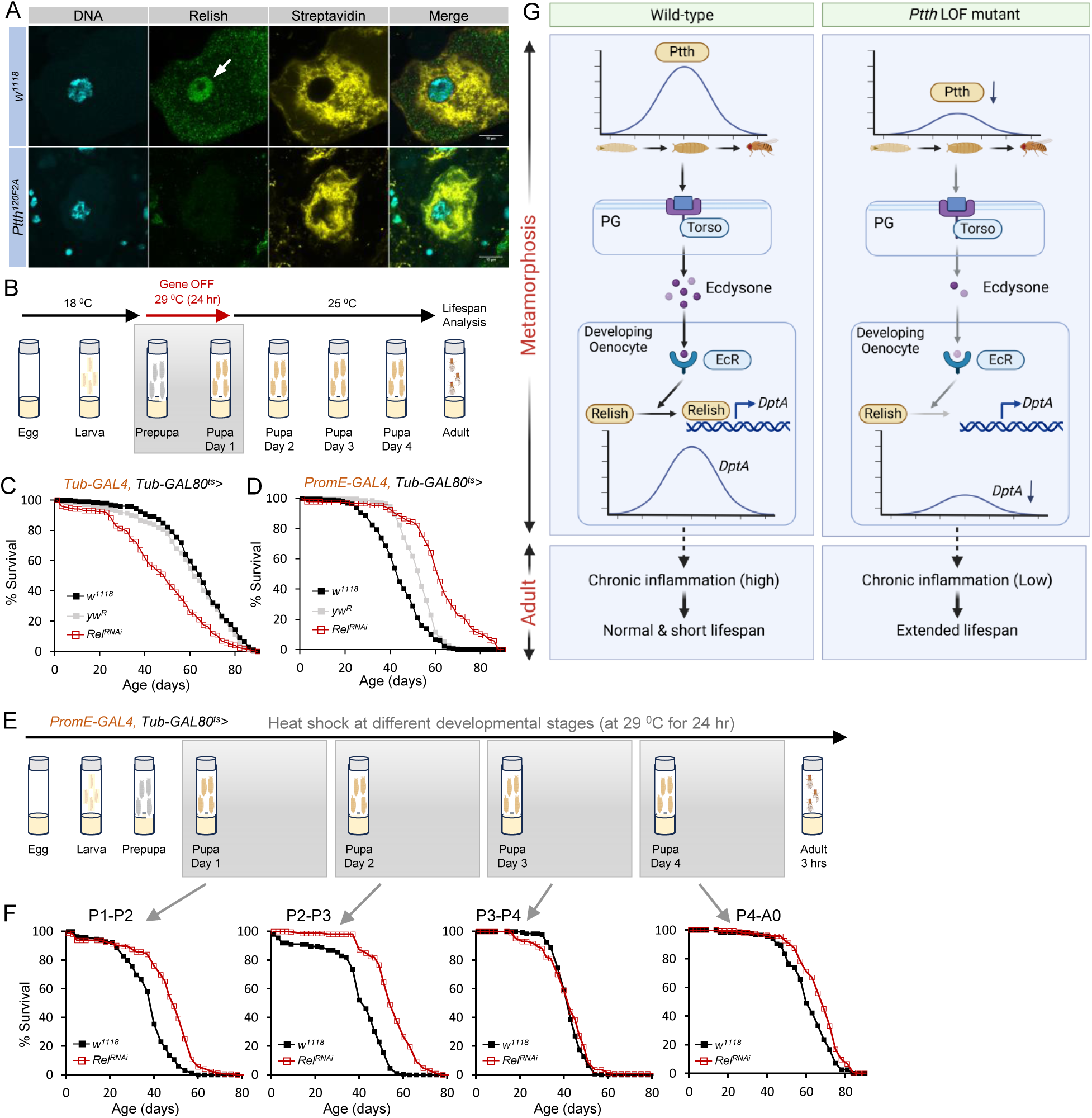
Pupal- and oenocyte-specific silencing of *Relish/NF-κB* prolongs lifespan. (*A*) Immunostaining analysis of nuclear translocation of Relish/NF-κB in oenocytes of day 1 pupa. Streptavidin staining was used to locate oenocytes. Scale bar: 10 μm. White arrow: nuclear localized Relish. (*B*) Schematic diagram showing the design of time-restricted gene silencing at early pupal stage. Flies were incubated at 29 ^0^C for 24 hours to activate Gal4 and RNAi (via inactivation of Gal80). (*C* and *D*) Lifespan analysis of whole body-(*Tub-GAL4; Tub-GAL80^ts^*) or oenocyte-specific (*PromE-GAL4; Tub-GAL80^ts^*) knockdown of *Relish* at early pupal stage (female). Two control flies were used, *yw^R^* and *w^1118^*. Log-rank test, *p* < 0.001, n = 553 (*Tub-GAL4; Tub-GAL80^ts^*) or 576 (*PromE-GAL4; Tub-GAL80^ts^*). (*E*) Schematic diagram showing the design of time-restricted gene silencing at various pupal stages. (*F*) Lifespan analysis of oenocyte-specific knockdown of *Relish* at different pupal stages (female). *w^1118^* was used control flies. P1: Log-rank test, *p* < 0.001, n = 344. P2: Log-rank test, *p* < 0.001, n = 414. P3: Log-rank test, *p* > 0.1, n = 332. P4: Log-rank test, *p* < 0.001, n = 280. (*G*) Proposed model to show insect hormone PTTH regulates lifespan through temporal and spatial activation of NF-κB signaling during *Drosophila* metamorphosis.

### Pupal- and oenocyte-specific silencing of *Relish/NF-κB* prolongs lifespan

As PTTH regulates NF-κB signaling in developing oenocytes, we wondered whether genetic manipulation of NF-κB signaling during metamorphosis would impact adult lifespan. We used the temperature-sensitive GAL80 system (*GAL80^ts^*) to achieve temporal and spatial gene silencing during *Drosophila* metamorphosis (**Fig. 6B**). Flies carrying *GAL80^ts^* and *GAL4* were maintained at 18 °C throughout larval development (*GAL4* expression is inhibited), and then switched to 29 °C at specific prepupal or pupal stages to activate *GAL4* expression for about 24 hours before switching to 25 °C for lifespan analysis (**Fig. 6B**). We first confirmed that the approach was efficient at silencing *Relish* during pupal development using *Tub-GAL4; Tub-GAL80^ts^*. As shown in **Fig S6A**, 24 hours of activation of *Relish* RNAi from white prepupa to one-day-old pupal stage resulted in a significant reduction in *Relish* expression at most pupal stages (up to 3-day-old pupal stage), while *Relish* expression was restored back to wild-type levels in adults. Surprisingly, global knockdown of *Relish* during pupal development significantly shortened lifespan (**Fig. 6C and S6B**), a finding consistent with a recent study showing that pupal-specific knockdown of *Relish* increases the susceptibility of adult flies to viral infection (35).

Since we found that PTTH regulates the activation of NF-κB signaling in developing oenocytes, we tested whether *Relish* knockdown in oenocytes during early pupal stages could extend the lifespan of adult flies. Strikingly, oenocyte-specific silencing of *Relish* during early pupal stages significantly prolonged adult lifespan (**Fig. 6D and S6C**). Given the peak expression of AMP genes during early pupal stages (P1-P2), we tested whether silencing *Relish* during early pupal stages (P1-P2) was required for lifespan extension. Indeed, when *Relish* was knocked down in oenocytes from P1 to P2 or from P2 to P3, but not from P3 to P4 or from P4 to A0, the lifespan was greatly prolonged (**Fig. 6E and 6F**). Altogether, our findings strongly suggest a novel temporal and spatial regulation of NF-κB signaling in developing oenocytes, which links animal development to adult lifespan.

## Discussion

Developmental signaling pathways are often involved in longevity and lifespan control, such as growth hormone (4, 7), insulin/IGF (5, 6), and mechanistic target of rapamycin complex 1 (mTORC1) signaling (43–46). However, it remains unclear how these signaling pathways link development to adult lifespan and when these programs are active during development. In this study, we uncovered a novel role for the insect hormone PTTH in lifespan regulation during *Drosophila* development. Specifically, we found that loss of *Ptth* prolongs lifespan by repressing age-dependent induction of NF-κB signaling and chronic inflammation in fly hepatocytes (oenocytes). Rather than targeting adult tissues, PTTH activates NF-κB signaling in developing oenocytes through Torso and EcR signaling during *Drosophila* metamorphosis. Strikingly, time-restricted and tissue-specific silencing of *Relish* in oenocytes during early pupal development significantly extends the lifespan of adult flies. Thus, our study unveils NF-κB signaling as a novel developmental program that is activated during the juvenile-to-adult transition, ultimately shaping adult physiology (**Fig. 6G**).

PTTH is well-known for its role in controlling the duration of larval growth and the development of adult tissues (15, 20). PTTH belongs to the cystine knot family of growth factors (17), and it has been proposed that PTTH functions as mammalian GnRH hormone in controlling the timing of the juvenile-to-adult transition (18, 19). In mammals, both GnRH and GH are involved in sexual maturation and puberty regulation. Genetic analysis on long-lived GH deficiency mice (e.g., Ames dwarf) suggests a mechanistic link between developmental programs and longevity regulation. This is further supported by the effect of early-life GH treatment on reversing the long lifespan of Ames dwarf mice (7, 47). Early-life GH treatment also promotes chronic inflammation late in life, highlighting the important role of GH in the regulation of inflammaging (7), which is reminiscent of what we observed in *Ptth* mutant flies. Although PTTH does not share sequence homology with mammalian GH, their roles in regulating the timing of juvenile development are similar. Besides, both PTTH and GH activate MAPK/ERK pathway in their target tissues (16, 48). Altogether, the findings from both flies and mice suggest an evolutionarily conserved mechanism by which growth factors (e.g., PTTH and GH) regulate developmental timing and adult lifespan.

Our studies suggest that PTTH may modulate NF-κB activation via ecdysone signaling during development and metamorphosis. However, it is known that ecdysone signaling functions to prime the target tissues for rapid immune activation upon infection (31), additional mechanisms are required to drive NF-κB activation during insect development and metamorphosis. During the prepupal and early pupal stages, the larval gut undergoes program cell death and autophagy-dependent degradation (49, 50). Thus, it is possible that during the larval gut breakdown, bacteria leak out of the gut to activate NF-κB signaling. On the other hand, activation of NF-κB might be due to intrinsic signals. For example, NF-κB can be activated in response to DNA damage through ATM (ataxia telangiectasia mutated) mediated signal transduction (51, 52). Further, cGAS-STING has emerged recently as a key regulator of antiviral immunity as it promotes NF-κB activation in response to sensing of cytosolic DNA (53, 54). Thus, DNA damage levels might be elevated during larval tissue destruction, which could lead to increased STING signaling and NF-κB activation. In support of this model, we observed an increased expression of *dSting* during the prepupal stage in our developmental RNA-seq analysis (data not shown).

Possibly, activation of NF-κB during metamorphosis could also reflect the remodeling of adult tissues. Interestingly, Dorsal (Dl), a REL domain-containing protein of the NF-κB family, was first identified as a regulator of dorsoventral pattern formation during *Drosophila* embryogenesis (55). NF-κB signaling has also shown to be activated in the hematopoietic niche to maintain blood progenitors in the developing lymph gland of *Drosophila* larvae (34). Also, during zebrafish development, NF-κB is activated in endothelial cells to drive the specification of hematopoietic stem and progenitor cells (56, 57). Further, NF-κB is required for TNF-α-mediated osteogenic differentiation from the human dental pulp stem cells, a type of mesenchymal stem cells (58). In addition, NF-κB activation promotes the migration and proliferation of human mesenchymal stem cells in response to proinflammatory cytokines, such as TNF-α and interleukin-1β (59, 60). Thus, it is possible that NF-κB signaling is activated by proinflammatory cytokines during *Drosophila* metamorphosis, while perturbation of NF-κB signaling might result in remodeling of adult tissues. Perturbation of NF-κB signaling in some tissues (e.g., oenocytes) could lead to protection against age-related damage at the adult stage.

In summary, we uncovered an unexpected activation of NF-κB signaling during *Drosophila* metamorphosis, which is under the control of PTTH. This temporal and spatial activation of NF-κB signaling is essential for oenocyte function and adult longevity. Our study provides novel insights into the developmental regulation of NF-κB signaling in shaping adult physiology, such as lifespan and healthspan. There are still many unanswered questions remaining. In particular, why and how fly hepatocytes (oenocytes) are protected from age-related damage by reducing NF-κB signaling during development.

## Materials and Methods

### Fly husbandry and stocks

Flies were maintained at 25°C, 60% relative humidity and 12-hour light/dark cycles. Adults were reared on agar-based diet with 0.8% cornmeal, 10% sugar, and 2.5% yeast (unless otherwise noted). Fly stocks used in the present study are: *Ptth^8BC1^* (generated by backcrossing *Ptth^8K1J^* with *w^1118^*), *Ptth^120BC2^ and Ptth^120BC3^* (generated by backcrossing *Ptth^120F2A^* with *w^1118^*), *Ptth^TI^* (BDSC #84568), *Da-GS-GAL4* (a gift from Marc Tatar), *Phm-GAL4*, *PromE-GAL4* (or *Desat1-GAL4.E800*, BDSC #65405), *PromE-GAL4, Tub-GAL80^ts^* (BDSC #65407), *Tub-GAL4, Tub-GAL80^ts^* (61), *UAS-Relish RNAi* (BDSC #28943), *UAS-FLAG-Rel.68* (BDSC #55777), *UAS-Ptth*-*RNAi* (VDRC #102043), *UAS-torso*-*RNAi (*VDRC #36280), *UAS-EcR*-*RNAi* (BDSC #29374). The following genotypes were used as control in the knockdown or overexpression experiments: *w^1118^* (20), *yw^R^* (a gift from Marc Tatar), *y^1^ v^1^; P[CaryP]attP40* (BDSC # 36304).

### Developmental timing analysis

To synchronize development for timed experiments, parental flies were allowed to lay eggs for 3∼4 hours on an apple juice agar plate coated with a thin layer of yeast paste. Twenty or twenty-four hours after egg laying, newly hatched L1 larvae were transferred to fly culture vials. Larvae were raised in groups of 30∼40 to prevent crowding. The time of pupariation was scored every 4 hours till all larvae molt into pupae. Pupariation data from 3 replicates were compiled and plotted in Excel or Graphpad.

### Demography and survival analysis

Flies were collected under brief CO_2_ anesthesia and placed in food vials at a density of 25∼30 females/males flies per vial, with a total of 150∼300 flies for most conditions. Flies were transferred to fresh food every other day, and dead flies were scored and counted. Survival analysis was conducted with JMP statistical software. Data from replicate vials were combined. Survival distributions were compared by Log-rank test.

### Climbing Assay

Climbing ability was measured via a negative geotaxis assay performed by tapping flies to the bottom of an empty glass vial and counting flies that climbed at different positions of the vial. Ten seconds after tapping, the percentage of flies in each section of the vial (0∼3 cm, 3∼6 cm, 6∼9 cm) were counted. The climbing ability index was calculated by weighing the number of the flies according to their positions of the test vial.

### Female fecundity analysis

Three-day-old mated female flies were maintained on food for 10 days at 5 females per vial and 3 vials per group. Flies were daily passed to new vials, and eggs were counted daily. The mean number of daily egg-laying was plotted.

### Oxidative stress resistance assay

To assess oxidative stress resistance, 5-day-old flies were transferred into glass vials containing 1% agar, 5% sugar, and 20 mM paraquat (Sigma, St. Louis, MO, USA). Dead flies were scored and counted every 4 hours. A total of 40∼50 flies were used for each genotype (10 flies per vial). Survival differences were analyzed by the Log-rank test.

### Bacterial challenge assay

Gram-negative pathogenic bacterium *Erwinia carotovora carotovora* 15 (*Ecc15*) was cultured overnight to obtain OD_600_ = 200. To assess the survival upon bacterial challenge, 5-day-old flies were infected with 1:1 mixture of 5% sucrose and 100X concentrated *Ecc15* overnight culture. The infection solution was added onto a filter disk that was placed over fly vials with 1% agar base. Dead flies were scored and counted every 4 hours. A total of 60∼80 flies were used for each genotype (10 flies per vial). Survival differences were analyzed by the Log-rank test.

### RNA extraction and Quantitative RT-PCR

Adult tissues (fat body, oenocyte, gut) were dissected in 1 × PBS before RNA extraction. For oenocyte dissection, we first removed the fat body through liposuction and then detached oenocytes from the cuticle using a small glass needle. Tissue lysis, RNA extraction, and cDNA synthesis were performed using Cells-to-CT Kit (Thermo Fisher Scientific). For whole-body RNA extraction, flies were collected on CO2 and transferred to a 1.7 ml centrifuge tube with a stainless steel ball and 500 μl Trizol reagent (Thermo Fisher Scientific, Waltham, MA, USA) and homogenized with Tissuelyzer. About 15 flies were used per replicate. DNase-treated total RNA was quantified by Nanodrop, and about 500 ng of total RNA was reverse transcribed to cDNA using iScript cDNA Synthesis Kit (Bio-Rad, Hercules, CA, USA).

QRT-PCR was performed with a Quantstudio 3 Real-Time PCR System and PowerUp SYBR Green Master Mix (Thermo Fisher Scientific). Two to three independent biological replicates were performed with two technical replicates. The mRNA abundance of each candidate gene was normalized to the expression of *RpL32* for fly samples by the comparative CT methods. Primer sequences are listed in the following: RpL32: forward 5’-AAGAAGCGCACCAAGCACTTCATC-3’ and reverse 5’-TCTGTTGTCGATACCCTTGGGCTT-3’. PGRP-LC: forward 5’-TTTAACCTTCCTGCTGGGTATC-3’ and reverse 5’-TTGTCTGTAATCGTCGTCATCTC-3’. DptA: forward 5’-TTGCCGTCGCCTTACTTT-3’ and reverse 5’-CCTGAAGATTGAGTGGGTACTG-3’. upd3: 5’-forward 5’-TCTGGAAGCTTCTTTCCGGC-3’ and reverse 5’-GCGGTCAGCTGTCGTCATTT-3’. Socs36E: forward 5’-ACTACGGTTTAGCCAAATTGC-3’ and reverse 5’-TGGACCTCCGATTGTTTTCTCT-3’. Relish: forward 5’-GAGCGTAATTGTGTCGAGGAA-3’ and reverse 5’-GGCAGATCCAGCGAGTTATTAG-3’.

### Immunostaining and imaging

To examine the nuclear translocation of Relish, adult oenocytes were dissected from one-day-old pupas or female flies in 1X PBS and then fixed in 4% paraformaldehyde for 15 min at room temperature. Tissues were washed with 1x PBS with 0.3% Triton X-100 (PBST) three times (∼5 min each time), and blocked in PBST with 5% normal goat serum for 30 min. Tissues were then incubated overnight at 4 °C with anti-Relish primary antibodies (RayBiotech RB-14-0004,1:500) diluted in PBST, followed by the incubation with secondary antibodies obtained from Jackson Immuno Research for 1 hr at room temperature the next day. After three washes, tissues were mounted using ProLong Gold antifade reagent (Thermo Fisher Scientific) and imaged with an FV3000 Confocal Laser Scanning Microscope (Olympus). DAPI or Hoechst 33342 was used for nuclear staining. Pupal oenocytes were marked with streptavidin Alexa Fluor 555 (Thermo Fisher Scientific).

### RNA-seq and bioinformatics

Two bulk RNA-seq analyses were performed separately to profile transcriptomic changes in two adult ages and eight developmental stages (see main text). Total RNA was collected from 10∼15 larvae, pupal or adult flies (three biological replicates each condition) using Trizol method (described as above), followed by DNase treatment (Ambion). RNA concentration was quantified by Qubit RNA BR Assay Kit (Thermo Fisher Scientific). RNA-Seq libraries were constructed using either NEBNext Ultra Directional RNA Library Prep Kit for Illumina (New England Biolabs) or by Novogene RNA-seq service. Poly(A) mRNA was isolated using NEBNext Oligo d(T)25 beads and fragmented into 200 nt in size. After first strand and second strand cDNA synthesis, each cDNA library was ligated with a NEBNext adaptor and barcoded with an adaptor-specific index. Libraries were pooled in equal concentrations and sequenced using Illumina HiSeq 3000 or Novoseq 6000 platforms.

The RNA-Seq data processing was performed on Ubuntu system. FastQC was first performed to check the sequencing read quality and Fastx is used to filter the bad quality read from fastq. Then the raw reads were mapped to the *D. melanogaster* genome (Drosophila_melanogaster.BDGP6.22.98.chr.gtf) using Star (https://github.com/alexdobin/STAR.git). Htseq-count was used to count the number of mapped reads on each gene and DE-seq2 (R package) was used to generate normalized data. After normalization, differentially expressed protein-coding transcripts were obtained using following cut-off values, false discovery rate (FDR) ≤ 0.05 and fold change ≥ 1.5 or 2. RNA-Seq read files have been deposited to NCBI ‘s Gene Expression Omnibus (GEO) (Accession # GSE271165 and #GSE271166). To review GEO accession GSE271165: Go to https://www.ncbi.nlm.nih.gov/geo/query/acc.cgi?acc=GSE271165. Enter token ynsdowwonhsnvcn into the box. To review GEO accession GSE271166: Go to https://www.ncbi.nlm.nih.gov/geo/query/acc.cgi?acc=GSE271166. Enter token crqrggyijrkljan into the box.

To identify stage- and genotype-specific differentially expressed genes from the developmental RNA-seq, the count matrix was prefiltered to remove low expressing genes in which the maximum expression level of a sample across all timepoints were less than 3 FPKM. Prefiltering also removes rows that have a total count that is less than 6. The filtered count matrix was then smoothed and normalized by the default method of DESeq2 (vs1.42). Differential expression analyses for stage-specific genes were performed using DESeq2 (one-tailed Wald test) between any one stage over the other seven stages for each genotype respectively. Differentially expressed genes (DEGs) were selected per timepoint as the genes having absolute log2 fold change (log2FC) larger than 1 and adjusted P value less than 0.05. To evaluate the difference between *Ptth* mutants and wild-type in each stage, we performed differential expression analysis (DESeq2, two-tailed Wald test) using filtered and smoothed counts matrices. In addition, multiWGCNA analysis was conducted to identify differentially expressed gene modules using a minimum module size of 50, maximum module size of 1000 and a soft threshold power of 12. All modules were tested for stage specificity (PERMANOVA *p*<10^−4^) given the genotype. Module genes are selected by choosing overlapping genes between corresponding wild-type and mutant modules along with visual inspection on heatmap.

### Statistical analysis

GraphPad Prism 7 (GraphPad Software, La Jolla, CA) was used for statistical analysis. To compare the mean value of treatment groups versus that of control, student t-test or one-way ANOVA (followed by Tukey’s multiple comparison) was performed. The effects of genotype on various traits were analyzed by two-way ANOVA followed by Bonferroni’s multiple comparison test.

## Acknowledgments

We thank the Bloomington *Drosophila* Stock Center (supported by NIH P40OD018537) and the Vienna *Drosophila* Research Center (VDRC) for fly stocks. We thank the *Drosophila* Genomics Resource Center (supported by NIH grant 2P40OD010949) for the cDNA clones. We thank FlyBase release (FB2024_02) for the data that was obtained and used in this study (62). We thank Drs. Naoki Yamanaka, Alex Gould, Marc Tatar, Pierre Leopold, Bowen Deng, and Yi Rao for providing fly stocks, fly information and suggestions. Graphical abstract and working model figures were created with BioRender.com. Work in the Perrimon lab is supported by 5P41GM132087 and 1U01AG086143. N.P. is an investigator of the Howard Hughes Medical Institute. This work was supported by National Institute on Aging AG058741 and AG075156 to H.B., Hevolution foundation HF-GRO-23-1199062-14 to H.B. and P.K.

---

**Fig S1.**
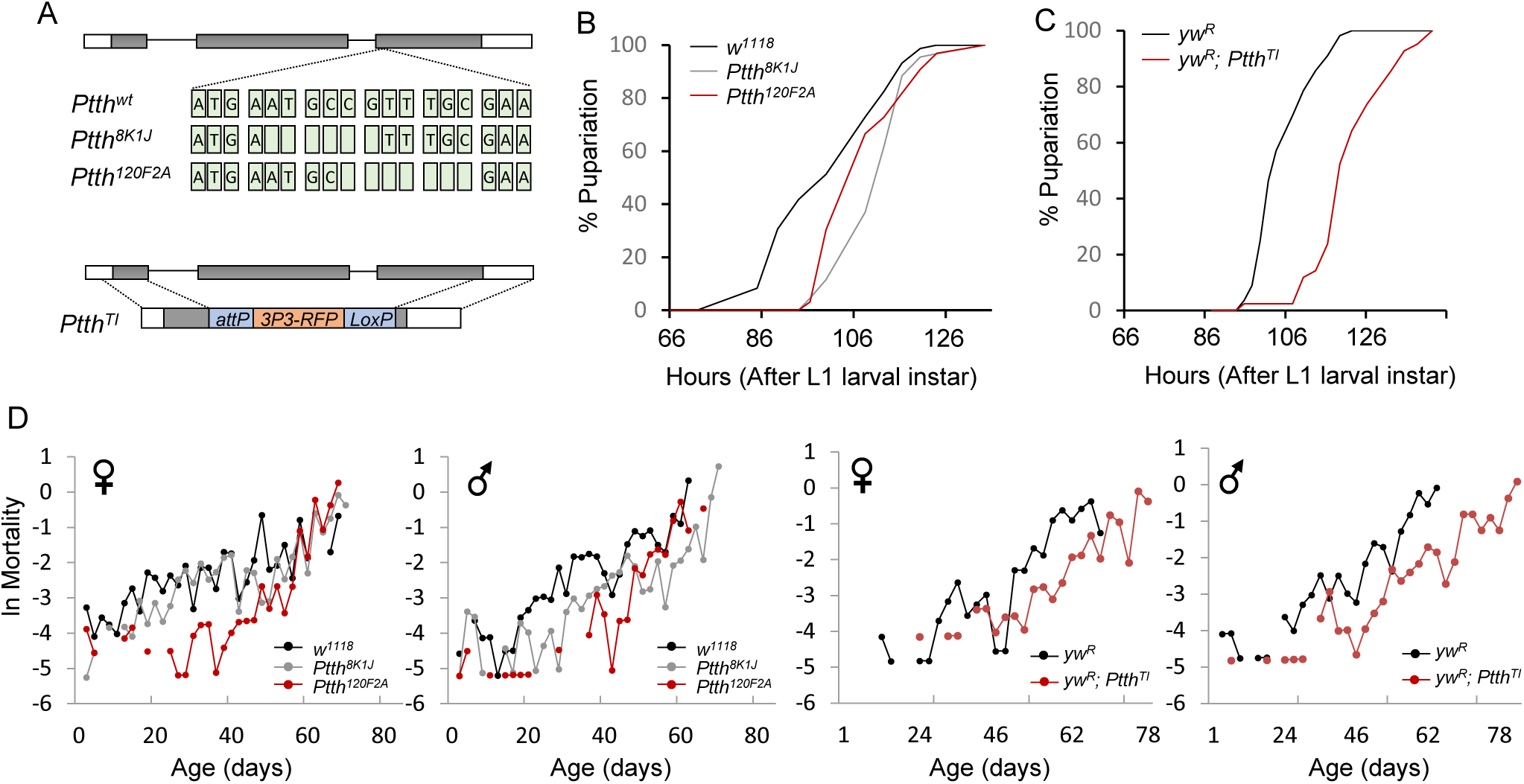
(*A*) Schematic diagram of three loss-of-function alleles of *Ptth*. (*B*) Developmental timing of wild-type (*w^1118^*) and *Ptth* mutants (*Ptth^8K1J^*, *Ptth^120F2A^*). Three replicates were performed for each genotype (about 30∼40 larvae each replicate). (*C*) Developmental timing of wild-type (*yw^R^*) and *Ptth* mutants (*Ptth^TI^*). (D) Mortality rate plots of wild-type and *Ptth* mutants. Mortality rate, ln(μ_x_), is calculated as ln(-ln(1-q_x_)), where q_x_ is age-specific mortality.

**Fig S2.**
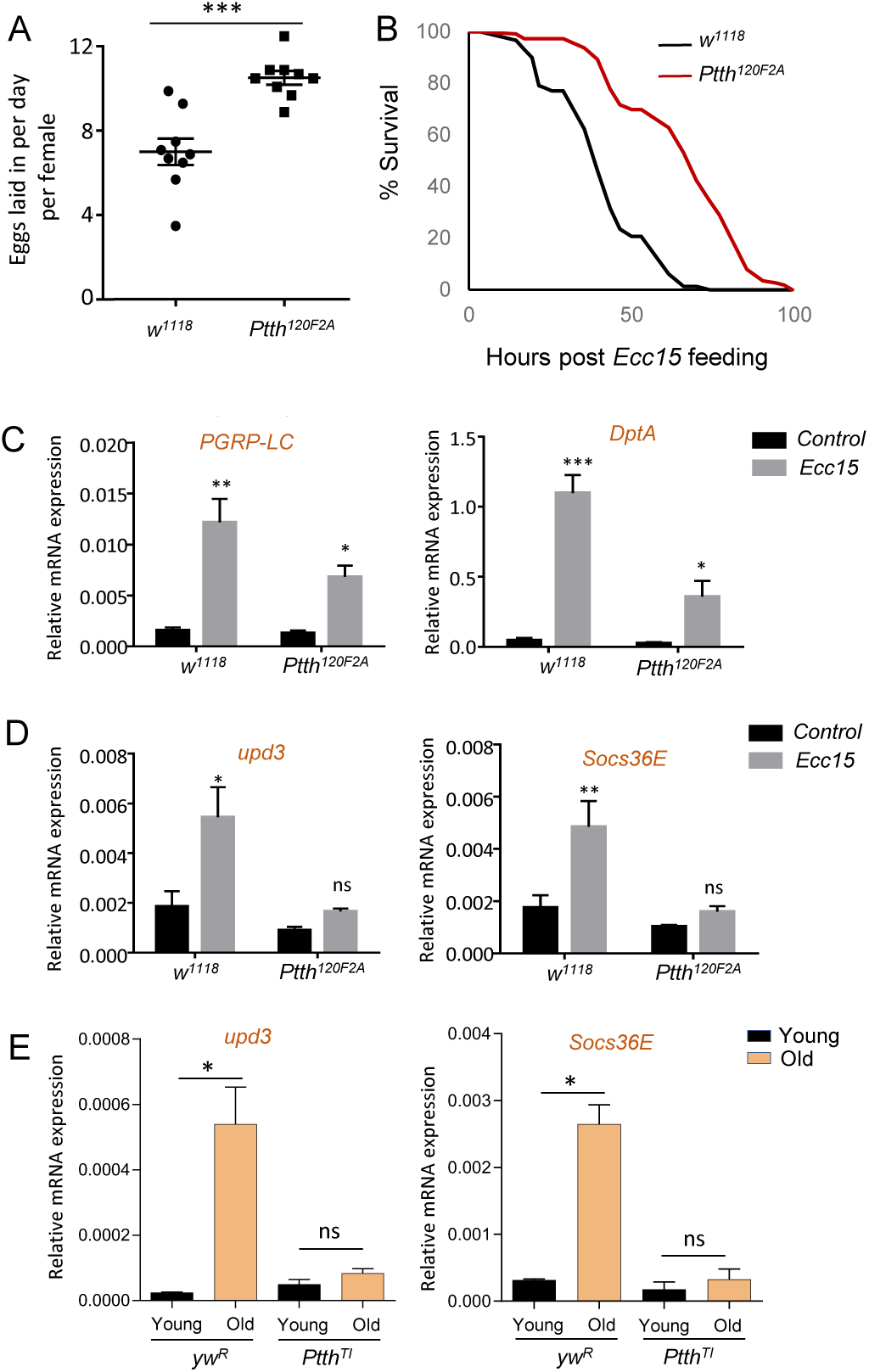
(*A*) Female fecundity analysis of young wild-type and *Ptth* mutants. Student t-test, *** *p* < 0.001, n = 9. (*B*) Survival analysis of young wild-type and *Ptth* mutants upon *Ecc15* treatment (female). Log-rank test, *p* < 0.001, n = 125. No mortality found in 5% sucrose control group. (*C* and *D*) The expression of *PGRP-LC, DptA, upd3*, and *Socs36E* of young wild-type and *Ptth* mutants upon 16 hours of *Ecc15* treatment (female). One-way ANOVA followed by Tukey’s multiple comparison test. ns, not significant; * *p* < 0.05; ** *p* < 0.01; *** *p* < 0.001. n = 3. (*E*) The expression of *upd3*, and *Socs36E* of young and old wild-type and *Ptth* mutants (female). One-way ANOVA followed by Tukey’s multiple comparison test. ns, not significant; * *p* < 0.05. n = 3.

**Fig S3.**
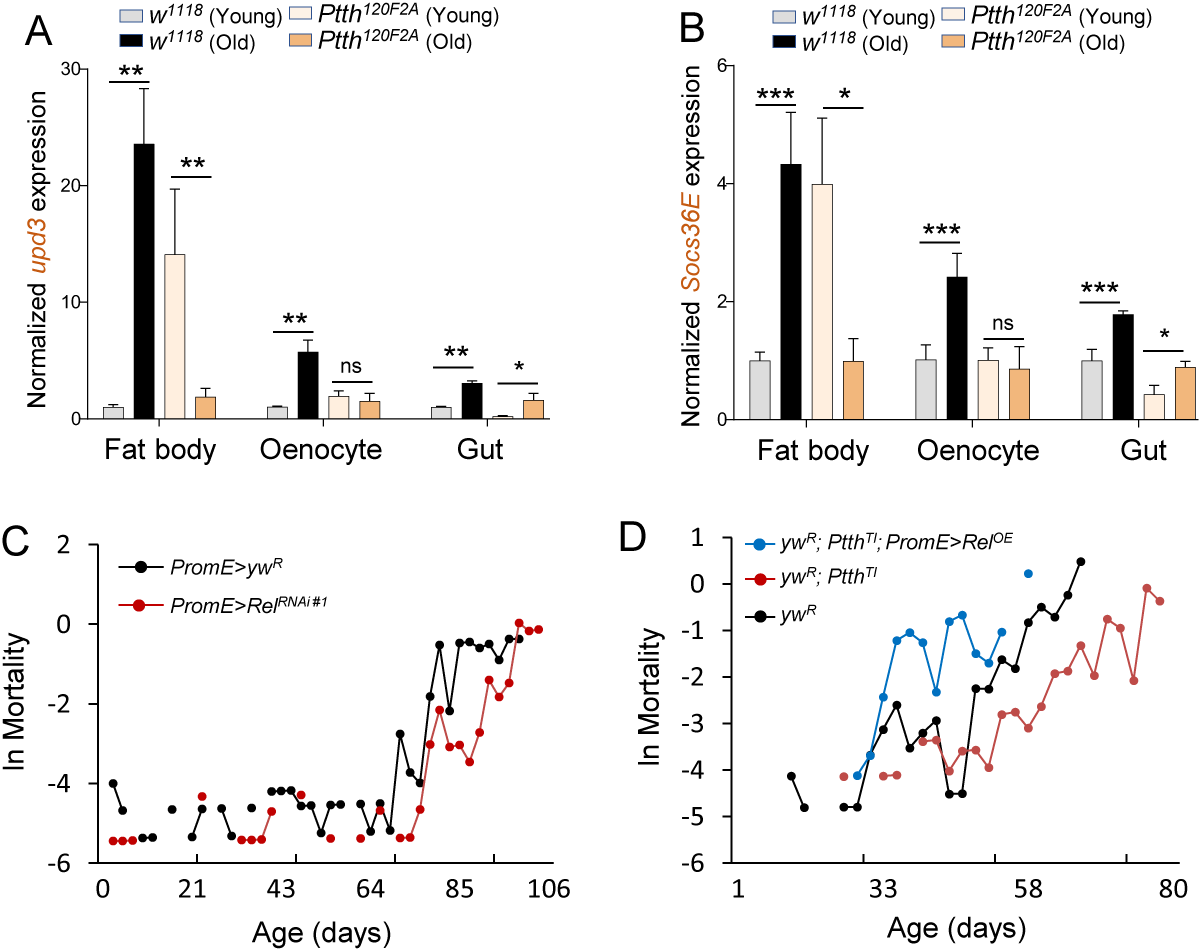
(*A* and *B*) qPCR analysis of the expression of *upd3* and *Socs36E* in three different fly tissues dissected from young and old *Ptth* mutants and wild-type (*w^1118^*). Two-way ANOVA followed by Bonferroni’s multiple comparison test. ns, not significant; * *p* < 0.05; ** *p* < 0.01; *** *p* < 0.001. n = 3∼6. (*C*) Mortality rate plots of oenocyte-specific *Relish* knockdown (female). Mortality rate, ln(μ_x_), is calculated as ln(-ln(1-q_x_)), where q_x_ is age-specific mortality. (*D*) Mortality rate plots of oenocyte-specific overexpression of *Relish/NF-κB* in *Ptth* mutant backgrounds (female).

**Fig S4.**
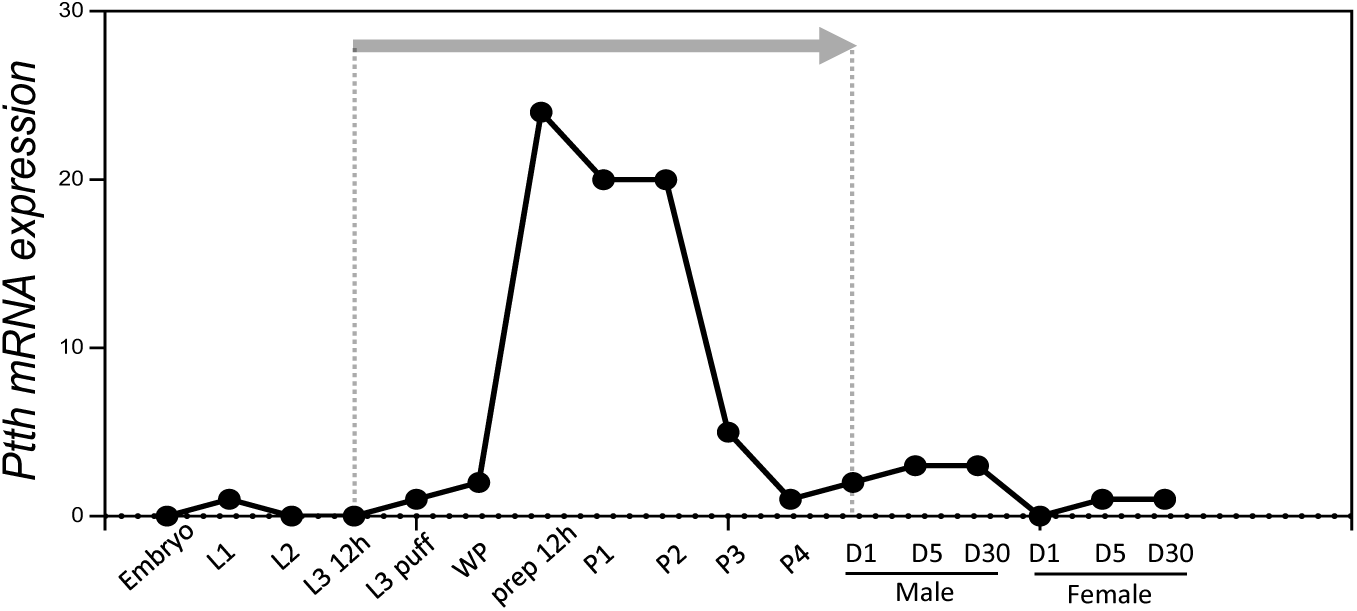
*Ptth* mRNA expression at different developmental stages. The expression value was retrieved from the FlyBase.

**Fig S5.**
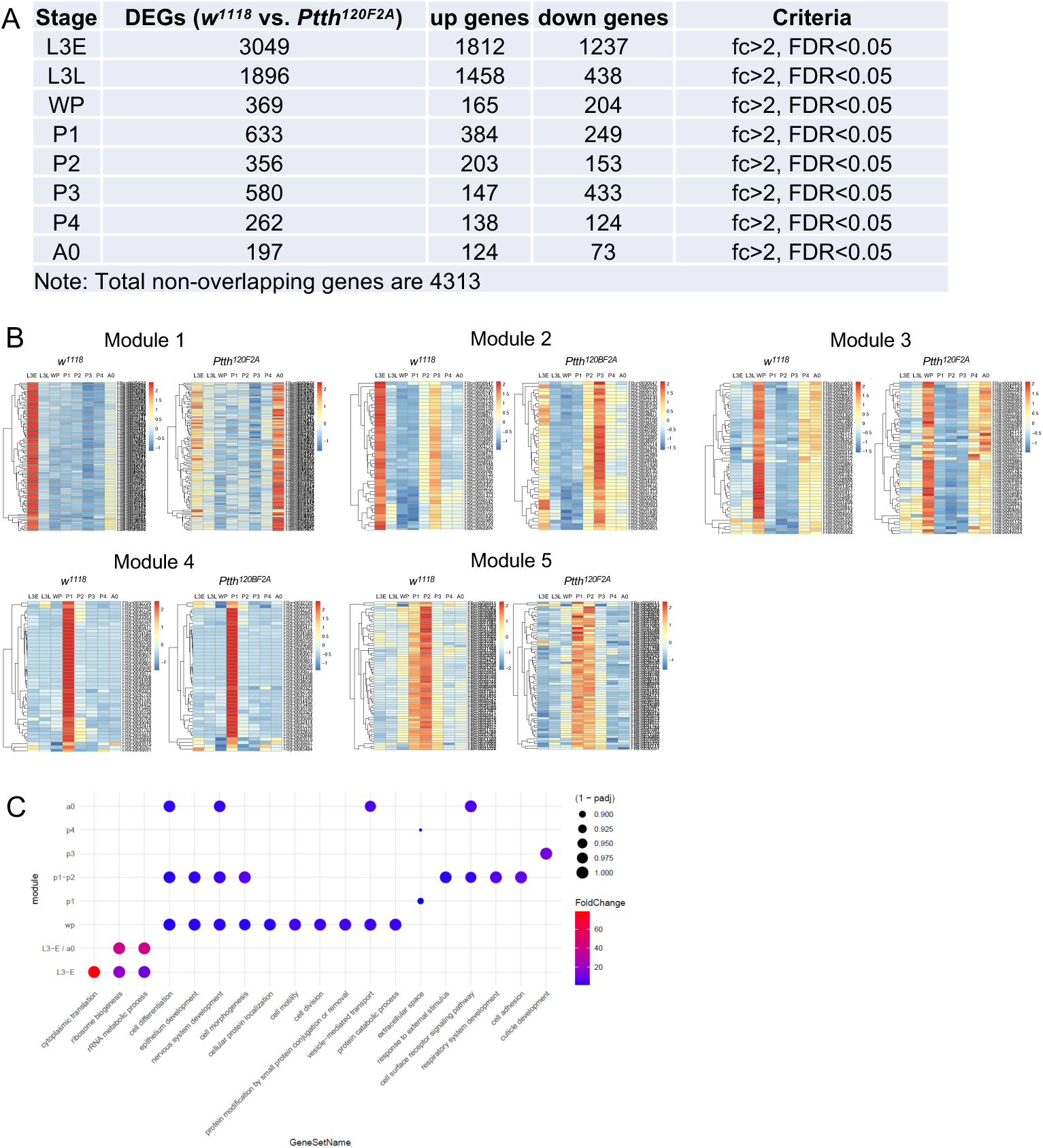
(*A*). The number of differentially expressed genes between wild-type and *Ptth* mutants across eight different developmental stages. (*B*) Five distinct modules identified by WGCNA analysis to show differentially expressed genes in wild-type (*w^1118^*) and *Ptth* mutants (*Ptth^120F2A^*) at different developmental stages. (*C*) GO term analysis for the biological processes enriched in different developmental stages.

**Fig S6.**
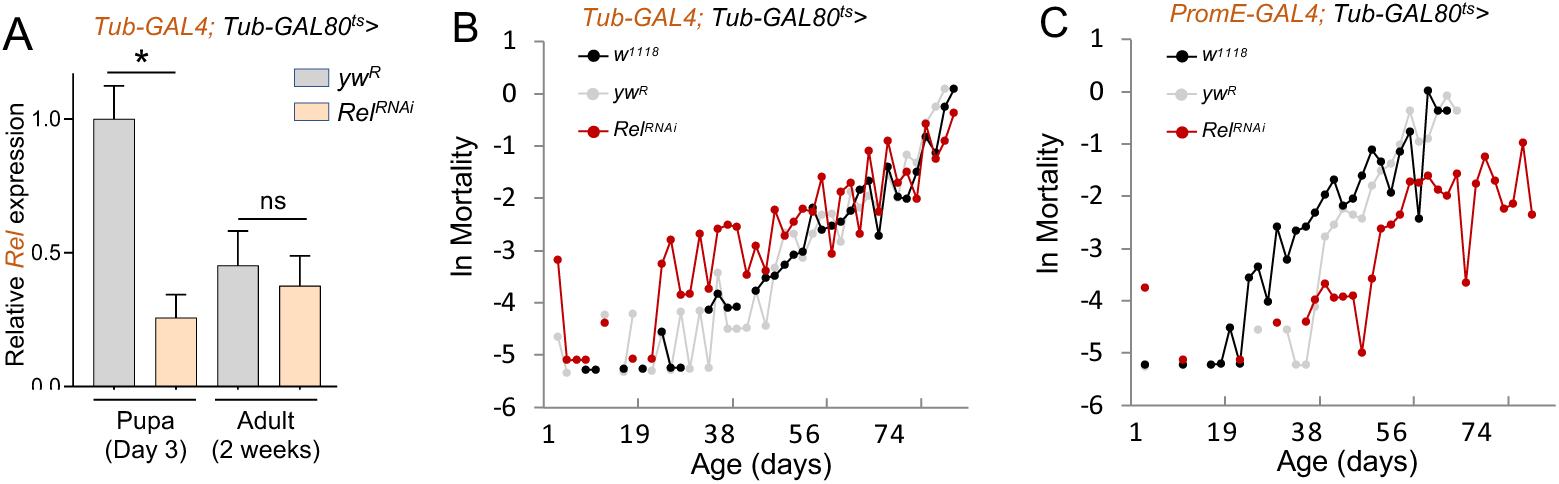
(*A*) qPCR analysis of the knockdown efficiency of *Relish* 2 days or 2 weeks post heat shock at 29 ^0^C. One-way ANOVA followed by Tukey’s multiple comparison test. ns, not significant; * *p* < 0.05. n = 3. (*B*) Mortality rate plots of pupal-specific whole body *Relish* knockdown (female). Mortality rate, ln(μ_x_), is calculated as ln(-ln(1-q_x_)), where q_x_ is age-specific mortality. (*C*) Mortality rate plots of pupal- and oenocyte-specific *Relish* knockdown (female).

